# From histology to macroscale function in the human amygdala

**DOI:** 10.1101/2024.07.09.602743

**Authors:** Hans Auer, Donna Gift Cabalo, Raul Rodriguez-Cruces, Oualid Benkarim, Casey Paquola, Jordan DeKraker, Yezhou Wang, Sofie Valk, Boris C. Bernhardt, Jessica Royer

**Author notes:** co-senior authors (Boris C. Bernhardt, Jessica Royer).

## Abstract

The amygdala is a subcortical region in the mesiotemporal lobe that plays a key role in emotional and sensory functions. Conventional neuroimaging experiments treat this structure as a single, uniform entity, but there is ample histological evidence for subregional heterogeneity in microstructure and function. The current study characterized subregional structure-function coupling in the human amygdala, integrating *post mortem* histology and *in vivo* MRI at ultrahigh fields. Core to our work was a novel neuroinformatics approach that leveraged multiscale texture analysis as well as non-linear dimensionality reduction techniques to identify salient dimensions of microstructural variation in a 3D *post mortem* histological reconstruction of the human amygdala. We observed two axes of subregional variation in this region, describing inferior-superior as well as medio-lateral trends in microstructural differentiation that in part recapitulated established atlases of amygdala subnuclei. Translating our approach to *in vivo* MRI data acquired at 7 Tesla, we could demonstrate generalizability of these spatial trends across 10 healthy adults. We then cross-referenced microstructural axes with functional blood-oxygen-level dependent (BOLD) signal analysis obtained during task-free conditions, and revealed a close association of structural axes with macroscale functional network embedding, notably the temporo-limbic, default mode, and sensory-motor networks. Our novel multiscale approach consolidates descriptions of amygdala anatomy and function obtained from histological and *in vivo* imaging techniques.

## INTRODUCTION

The amygdala is a central hub for socio-affective and cognitive functioning (1–3). Over the past decades, lesion studies in animals and humans have been crucial in our understanding of this structure’s functional role. Studies performed in animal models have reported significant deficits in a vast array of social and affective functions following amygdala lesions, including affective blunting, perturbed social interest and affiliation behaviors, increased aggression, altered sexual and maternal behaviors as well as fear response to environmental stimuli (4–6). In addition to altering social behavior, lesions of this structure in humans have been associated with impaired decision making (7) and deficits in attention and arousal mechanisms (8), emphasizing the importance of the amygdala in a broad array of functional domains.

Although earlier work on amygdala function in humans has considered this region as a single, unitary structure, the distinct roles of its individual subdivisions are now increasingly highlighted (9–11). Notably, early research on the amygdala in non-human primates has been instrumental in understanding its intricate structure, function and patterns of anatomical connectivity (12,13). This foundational study highlights the amygdala’s different subdivisions, most notably the basomedial nucleus (BM), basolateral nucleus (BL), and central nucleus (Ce) (14). Furthermore, this work describes a dense network between these subdivisions and the prefrontal cortex, most strongly found in the posterior orbitofrontal and anterior cingulate areas.

In humans, qualitative examinations of *post mortem* specimens have identified several subdivisions within the amygdala, each with distinct cytoarchitectural characteristics and distinguishable connectivity profiles (15). These individual subnuclei have often been grouped into larger subdivisions, specifically centromedian, laterobasal and superficial regions (16). Distinguishable connectivity profiles in these subdivisions have been previously observed through the analysis of resting-state functional magnetic resonance imaging (rsfMRI) (13,17). This non-invasive technique has been instrumental to interrogate grey matter (GM) connectivity and map functional networks in the brain by detecting coordinated hemodynamic signal fluctuations across regions (18–23). For instance, previous work has shown synchronized functional signals between the centromedial (CM) subdivision of the amygdala and middle and anterior cingulate cortices, frontal cortex, striatum, insula, cerebellum and precuneus, supporting processes such as attention control and visceral responses (24–26). Conversely, the laterobasal (LB) region shows unique connectivity with the inferior and middle temporal gyri and middle occipital gyrus which have been associated with associative processing of environmental information and the integration with self-relevant cognition for decision making (24,27–29). The superficial (SF) subdivision of the amygdala has rather been associated with social information processing and social interaction via its unique connectivity to the paracentral lobule, posterior cingulate cortex, and orbitofrontal cortex (17,30–33). Given these differences across amygdala subregions, combining structural and functional analyses can shed light on the multi-faceted contribution of the amygdala to affective and cognitive functioning by potentially revealing variable participation of its subdivisions in different functional networks.

Our current understanding of the amygdala highlights its multidimensional roles, supported by its complex anatomy and participation in multiple brain networks. However, microstructural atlases of this area developed using quantitative techniques are still lacking but are essential for large-scale investigations of structure-function coupling within this region. Emerging strategies for quantitative segmentations of amygdala subdivisions have shown promising results. For example, a dual-branch convolutional network model trained with features extracted from T1- weighted images could find strong overlap between automatically segmented labels and a manual segmentation of lateral, basal, cortico-superficial and centromedial subregions (34). Similarly, a machine learning-based correction model could achieve comparable accuracy using a multi-atlas segmentation model (35). Despite the promise of these methods, they remain to be validated in histology (36), which remains a key technique to validate MRI-based features with access to ground-truth measures of cytoarchitecture (37–39). Indeed, regions with different cytoarchitecture often show distinct myelination patterns, which can be observed through various MRI contrasts. Such myelin-sensitive imaging contrasts can differentiate regions with distinct intracortical myeloarchitectonic profiles, demonstrating ties between variations in cellular architecture and myelin distribution (40,41).

Our study seeks to elucidate the intricate structure-function relationships within the amygdala by leveraging advanced data-driven quantitative methods and high-resolution histology. Our approach first leveraged computer vision techniques to map major subdivisions of the amygdala in BigBrain (42), a *post mortem*, high-resolution 3D histological dataset providing direct measurements of brain cytoarchitecture. We translated this approach to *in vivo* MRI data acquired at ultra-high fields enabling individual-specific assessments of microstructure-function coupling in the amygdala. Harnessing myelin-sensitive contrasts and multi-echo rsfMRI, our study identifies a principal axis of microstructural and functional network dissociation within the human amygdala from a data-driven analysis of its cyto-and myeloarchitecture.

## RESULTS

### Data-driven histological analysis of the human amygdala

Using a radiomics approach (43,44), we computed the four central moments (mean, variance, skewness, and kurtosis) from cell body staining intensities in the amygdala provided by the BigBrain dataset (42,45,46), **Figure 1A**). These voxel-wise metrics were computed across different kernel sizes, with local 3-dimensional neighborhoods ranging from a radius of 2 to 10 voxels. As expected, increasing kernel values produced smoother images for all central moments. Thus fine-grain features were emphasized at small kernel values and coarser features with larger kernel values.

**Figure 1.**
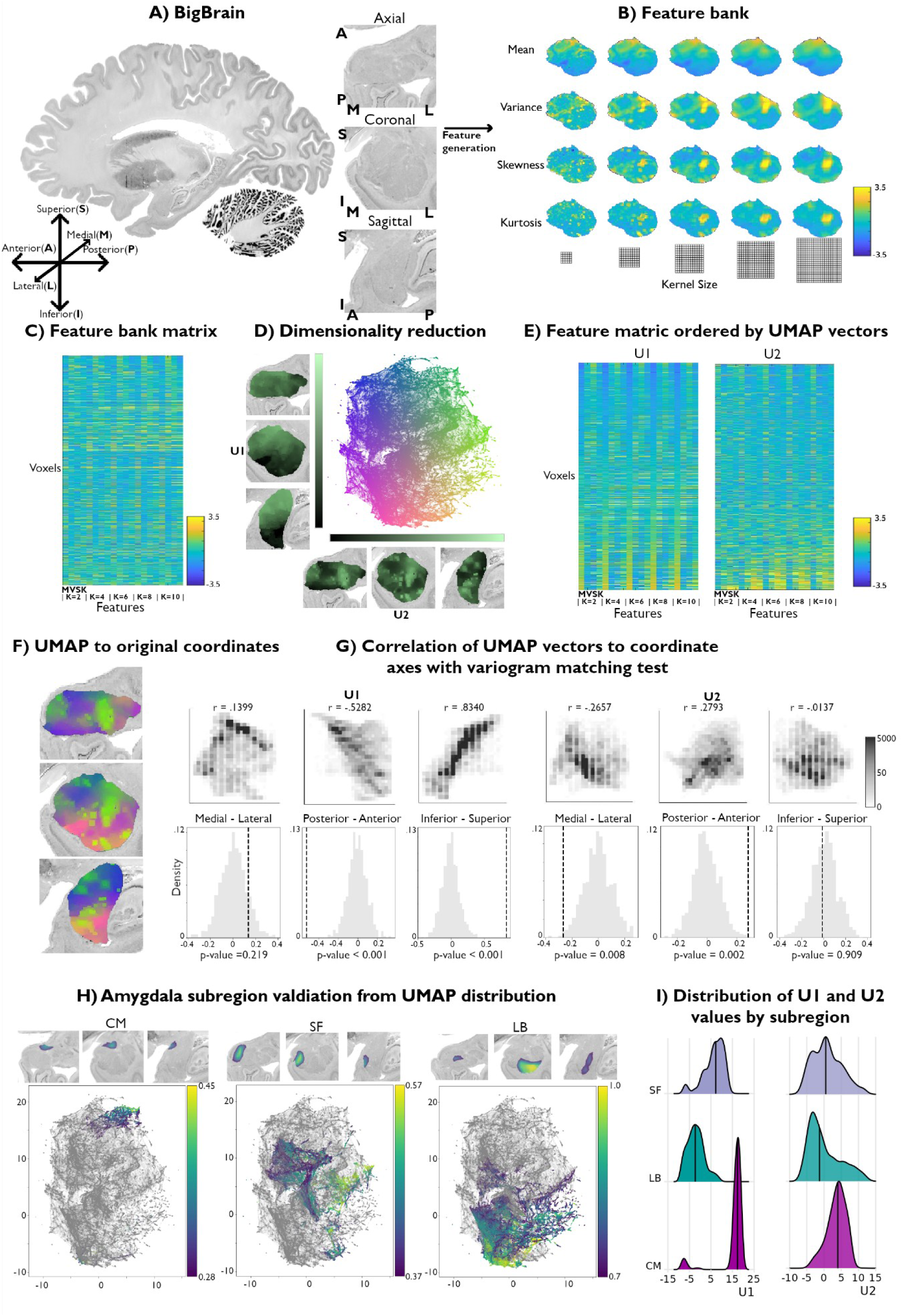
Data-driven histological mapping of the human amygdala. **(A)** The amygdala was segmented from the 100-micron resolution BigBrain dataset using an existing subcortical parcellation (45). Slice orientation of subpanels containing amygdala images is consistent across all panels in this figure. **(B)** Leveraging the pyRadiomics package v3.0.1 (47), we built a multiscale histological feature bank of the amygdala capturing fine-to-coarse intensity variations within this structure. Feature values were all normalized to better visualize relative intensity differences. **(C)** Matrix representation of the normalized feature bank shown in *A*. **(D)** We applied UMAP to this feature bank to derive a low dimensional embedding of amygdala cytoarchitecture, defining a 2-dimensional coordinate space (scatter plot, middle). Colors of the scatter plot represent proximity to axis limits. **(E)** Reordering the feature bank according to each eigenvector (U1 and U2) highlights the underlying variance in each feature captured by UMAP. **(F)** Coloring each amygdala voxel according to its corresponding location in the UMAP embedding space partially recovered its anatomical organization. **(G)** U1 and U2 were correlated to the three spatial axes and variogram matching tests assessed the statistical significance of each correlation. **(H)** Coloring the embedding space with openly available probabilistic map labels of the three main amygdala subregions showed that this region’s microstructural architecture could be recovered by UMAP. **(I)** Ridge plots of the probability values per subregion also illustrate a characterization of the subregions in U1.

The resulting feature bank showed heterogeneous feature profiles across selected moments and kernel sizes (**Figure 1B**). Mean intensities smoothly increased in the ventral to dorsal direction, culminating in highest intensity values in the dorsal subnucleus regions, and mainly coinciding with the CM subdivision of the amygdala. Variance, however, was highest along the amygdala’s borders with the entorhinal cortex and in the amygdala-striatum transition zone. Skewness and kurtosis maps generally co-varied spatially and both highlight high skewness and kurtosis within the lateral nucleus. Together, these findings indicate that the selected features captured both unique and shared characteristics of histological signal variations within the amygdala.

We then applied Uniform Manifold Approximation and Projection (UMAP), a non-linear dimensionality reduction technique that preserves the local and global structure of high-dimensional data by projecting it into a lower-dimensional space (48) to the resulting 20-feature matrix (**Figure 1C**) to derive a 2-dimensional embedding of amygdala cytoarchitecture (**Figure 1D**). This approach allowed us to bring the high dimensional histological feature space to a 2D embedding space composed of every amygdala voxel. As such, amygdala voxels were ordered along two dimensions, U1 and U2, capturing two axes of variance in amygdala cytoarchitecture. To better visualize which features may be driving each UMAP dimension, we sorted all input features along U1 and U2 (**Figure 1E**). Ordering the mean along U1 highlighted increasing intensity values at all different kernel sizes, where highest intensity voxels co-localized with low values in U1. While the variance, skewness and kurtosis showed less systematic changes at lower kernel values, patterns became more evident at higher kernels. Indeed, variance and kurtosis seemed to show an opposite trend to the mean along U1, where higher skewness and kurtosis co-localized more strongly with positive values of U1 (**Supplementary Table S1.1**). In contrast, sorting the feature matrix by U2 showed very similar trends between all features, independent of its moment and kernel value. More specifically, the highest intensity voxels were mostly found to show higher values along U2 (**Supplementary Table S1.2**). Overall, these UMAP-driven visualizations of the histological feature space suggest our dimensionality reduction approach could recover moment-specific (U1) as well as global intensity covariations across moments (U2).

To contextualize variations in U1 and U2 values across the amygdala, we computed voxel-wise correlations between each UMAP component and the amygdala coordinate space. U1 primarily varied along the inferior-superior axis (U1: r = 0.8340; U2: r =-0.0137), followed by posterior-anterior (U1: r =-0.5282; U2: r = 0.2793) and medial-lateral directions (U1: r = 0.1399; U2: r =- 0.2657) (**Figure 1G**). Statistical significance of correlations was assessed using a variogram matching approach (49) implemented in the BrainSpace toolbox (50) (**Figure 1G**). Both UMAP components were found to significantly co-vary along the posterior-anterior axis (U1: *p_null_*< 0.001; U2: *p_null_* = 0.002), while only U1 was significantly correlated with the inferior-superior axis (U1: *p_null_* < 0.001; U2: *p_null_* = 0.909). Neither U1 or U2 were significantly correlated with the medial-lateral coordinate (U1: *p_null_* = 0.441; U2: *p_null_* = 0.068).

### Validation of histological space using independent post mortem dataset

We contextualize our data-driven approach using established probability maps of amygdala microstructure. These openly available probability maps generated from visual inspection of 10 *post mortem* brains divide the amygdala into CM, SF, and LB subregions (39). Maps were thresholded to retain only voxels with highest 5% probability values and binarized. Plotting the probability values of retained voxels for each subregion showed that UMAP could dissociate these established cytoarchitectural subdivisions of the amygdala (**Figure 1H**). The three subregions could be particularly segregated along the U1 component. Indeed, we found significant differences in U1 values across the three amygdala subdivisions (F = 1630.8; *p_null_*< 0.001), while no significant difference was found across U2 (F = 30.3; *p_null_* < 0.581) (**Figure 1I**). Together, these findings show our theoretically grounded framework can successfully distinguish different subregions in the amygdala in a purely data-driven way. Additionally, results could be replicated when analyzing signals from the right amygdala, supporting the potential generalizability of our framework (**Supplementary Figure S1**).

### In vivo generalizability of histological space

We also assess the generalizability of these results to *in vivo* myelin-sensitive MRI data. We leverage quantitative T1 imaging collected at a field strength of 7 Tesla (7T) in 10 unrelated, healthy participants (**Figure 2A**). These images offered a resolution of 500um and were run through a similar analytical framework as the BigBrain dataset. We used individual-specific segmentations of the amygdala obtained with VolBrain (51). Once again, a feature bank was rendered from the same central moments, specifically mean, variance, skewness, and kurtosis. Kernel sizes varied from size 1-5, resulting in 20 distinct feature maps (**Figure 2A**). The new feature bank was again submitted to UMAP for dimensionality reduction (**Figure 2A**), and values of each component were plotted back to their respective coordinates in the amygdala.

**Figure 2.**
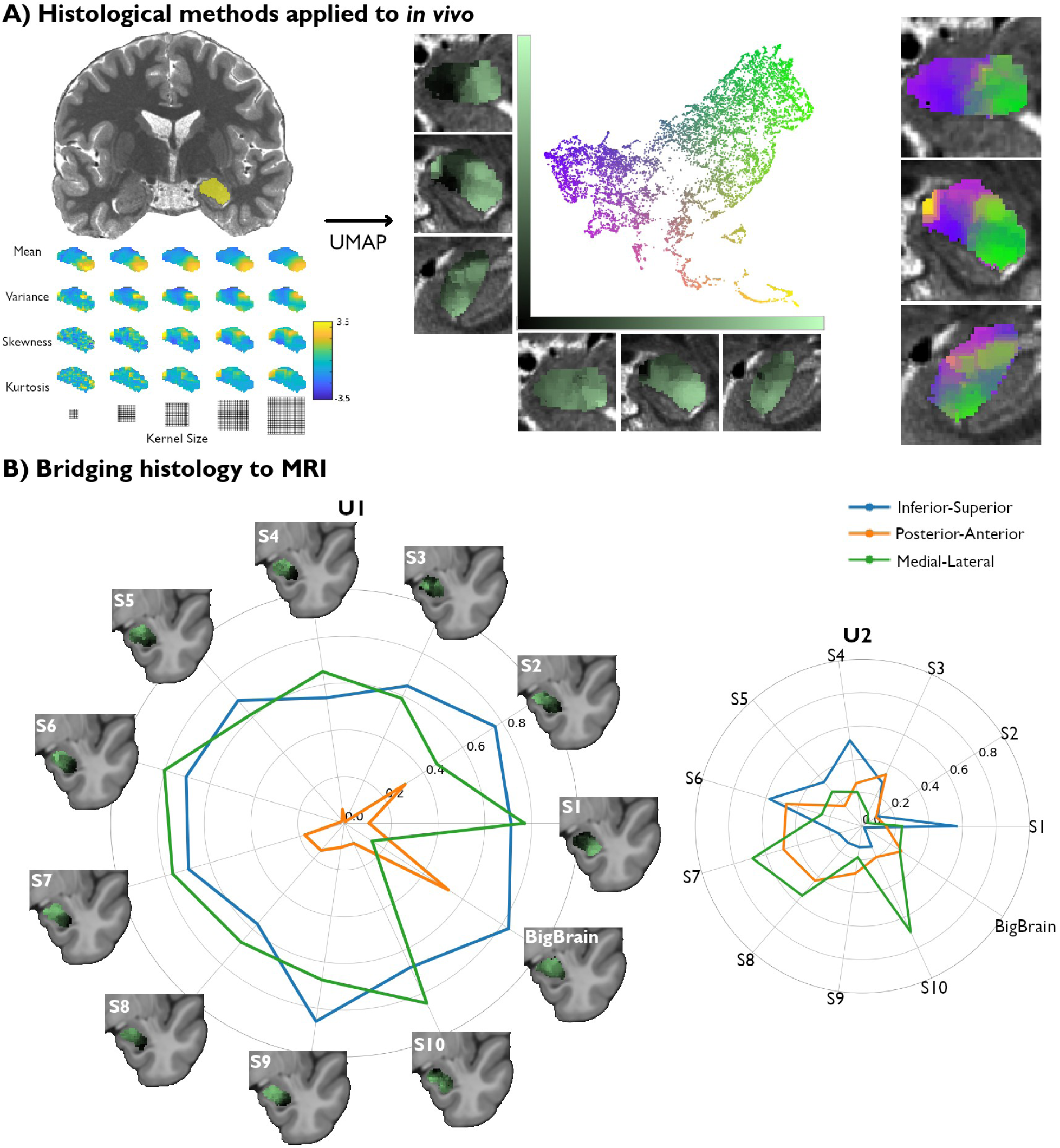
Translating amygdala histological space to *in vivo*, ultra high-resolution, myelin-sensitive MRI. **(A)** We segmented the left and right amygdalae of individual subjects from quantitative T1 (qT1) scans, and applied the same framework as developed in *post mortem* imaging to derive subject-specific, *in vivo* representations of amygdala microstructure. **(B)** Correlation values between the UMAP components (U1 and U2) and the three coordinate axes of the 10 MRI subjects were computed in MNI152 space and then contrasted with the correlation values found in the histological data (BigBrain transformed to MNI152 space).

We then compared spatial variations of the two UMAP components uncovered in each participant with the UMAP space derived from the BigBrain dataset. When examining the spatial layout of the UMAP components, we find similar neuroanatomical trends in all subjects to those found in histology (**Figure 2B**). Notably, the U1 vector of all subjects are found to be significantly correlated to the inferior-superior axis from a variogram matching test (**Supplementary Table S2.1**), similarly to BigBrain, suggesting the potential for this framework to capture important structural features in histology as well as *in vivo* MRI. We also find that the medial-lateral axis correlations to U1 across all subjects are consistent from the variogram test (**Supplementary Table S2.1**). The spatial layout of U2, on the other hand, showed lower consistency across participants, as none of the coordinate axes were significantly correlated with U2 in more than 7/10 subjects (**Supplementary Table S2.2**). In sum, we identify a single axis (U1) able to pick up on important amygdala microstructural features in both *post mortem* histology and *in vivo* markers of GM microstructure.

### Association with macroscale function

We next sought to investigate associations between amygdala microstructural organization and this region’s macroscale functional organization across subjects. Given U1’s strong spatial consistency across participants and microstructural modalities, we generated subject-specific masks segregating this component’s highest and lowest 25% of values within the amygdala for each participant. This resulted in two distinct regions of interest, reflecting the anchors of maximal microstructural dissociation within the amygdala for each participant, from which we could extract corresponding functional activity recorded at rest (**Figure 3A**).

**Figure 3.**
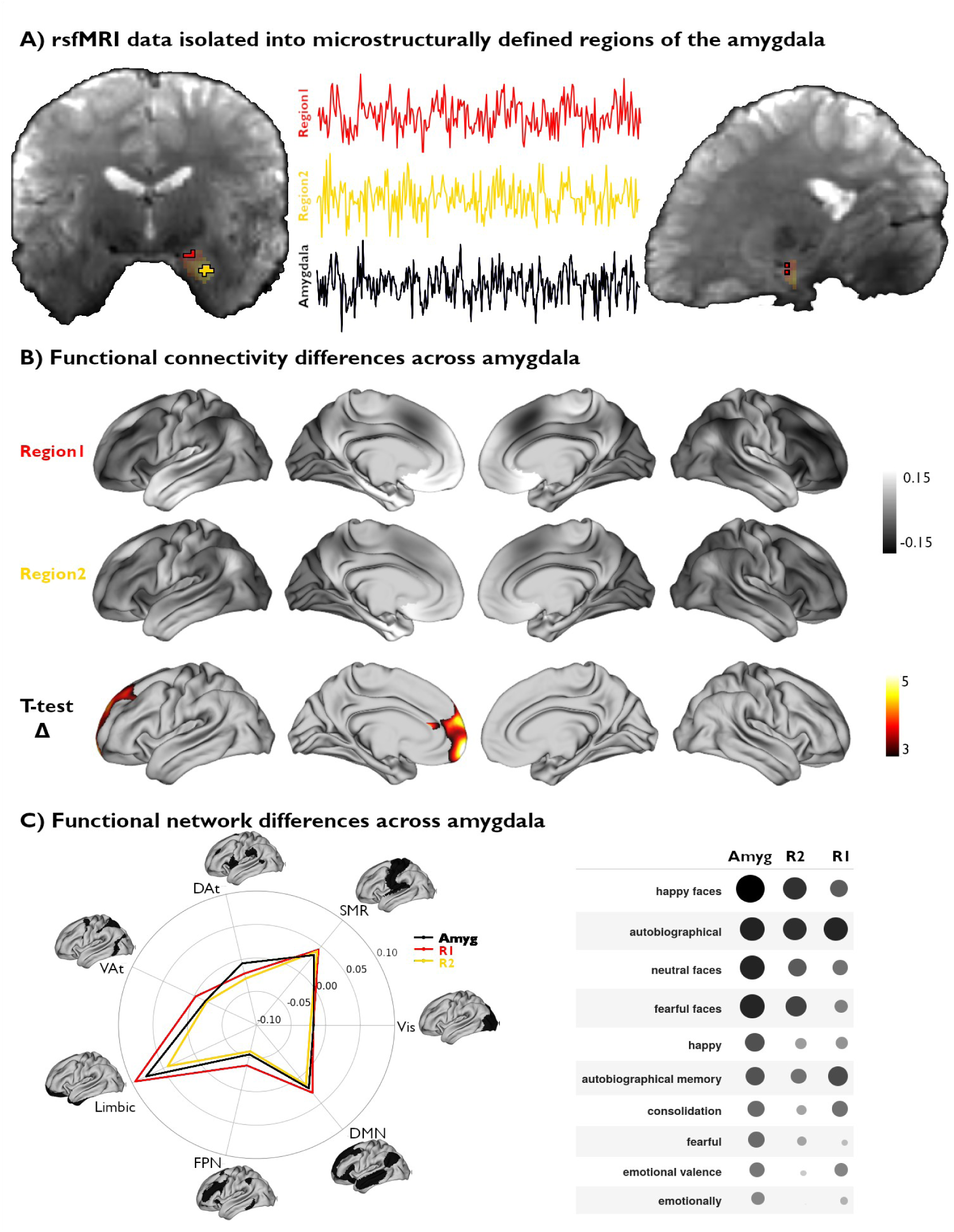
Functional network mapping of amygdala microstructural subregions. **(A)** We isolated the rsfMRI timeseries of two amygdala subregions, defined from subject-specific U1 topography, as well as the whole amygdala. **(B)** We computed the functional connectivity of both amygdala subregions, and project resulting correlations to the cortex. We further demonstrate the differences in connectivity patterns between both subregions (t-value) and highlight the regions with significant differences (*pFWE*<0.05). **(C)** Left: The activation patterns illustrated in (B, top) were averaged within intrinsic functional communities defined by Yeo, Krienen, et al. (2011). Right: Meta-analytic decoding of functional connectivity patterns of both amygdala subregions and the whole amygdala dissociated cognitive and affective functional affiliations of this region.

Voxel-wise rsfMRI timeseries were averaged within each microstructural subregion and correlated to vertex-wise cortical timeseries. A linear mixed effect model comparing amygdalo-cortical connectivity profiles between both subregions showed that functional network affiliations significantly differed across U1 subregions, with stronger connectivity observed between the superior portions of the amygdala (top 25% U1 values) and the prefrontal lobe (**Figure 3B**). Stratifying functional connectivity patterns of each amygdala subregion to the cortex according to established intrinsic functional network communities further highlighted the relatively stronger connectivity of the superior subregions to all cortical networks, particularly the limbic, frontoparietal, and default mode networks (**Figure 3C**). Meta-analytical decoding of subregional connectivity profiles using NeuroSynth (52) emphasized the functional dissociation in connectivity patterns of both microstructurally-defined areas. This analysis showed that the functional connectivity pattern of the region with highest 25% U1 values was most strongly associated with terms relating to autobiographical memory (‘autobiographical’ and ‘autobiographical memory’), while the other seed region’s connectivity profile overlapped with activation patterns related to emotional input (‘happy faces’, ‘neutral faces’ and ‘fearful faces’) (**Figure 3C**). Furthermore, decoding our statistical effects map (Region 1 connectivity > Region 2 connectivity) highlighted associations with terms relating to the self, introspection, and reward (‘self referential’, ‘referential’, ‘moral’, ‘autobiographical’, ‘smoking’, ‘craving’). Collectively, these findings show that our theoretically-grounded approach, developed in histology and generalizable to microstructurally-sensitive *in vivo* MRI data, can delineate distinct functional network embeddings in the human amygdala.

## DISCUSSION

The amygdala is a crucial structure for several aspects of cognitive and socio-affective functioning (3). These functions are supported by complex connectivity patterns to other brain regions, stemming from distinct subnuclei with unique microstructural properties (15). However, current investigations of structure-function coupling in the amygdala are limited by a lack of datasets and tools for individualized and observer-independent delineation of its subregions. Indeed, the strong inter-individual variability of its structural and functional organization (16,53) motivates more personalized approaches to reliably study the microstructural determinants of amygdala function and connectivity. In the current paper, we present a data-driven exploration of subcortical cytoarchitecture applied to the human amygdala. We could translate this approach to microstructurally-sensitive *in vivo* MRI data as a bridge, to ultimately examine associations between microstructural subregions and functional networks. As such, the present work defines a quantitative and integrated account of the amygdala’s microstructural composition and functional organization. In doing so, our approach sets the stage for novel investigations spanning other subcortico-cortical systems, and shows potential to deliver new insights into brain-wide principles of structure-function coupling and how this interplay may be altered in clinical populations.

The proposed framework aimed to delineate amygdala subnuclear organization by leveraging a multiscale texture processing pipeline designed to retain finer and coarser regional cytoarchitectonic properties. We specifically harness radiomics, a field with established diagnostic and prognostic potential in medical imaging (54). Feature selection in our study was motivated by previous work conducted at the level of the neocortex (36,43) and focused on the four central moments, specifically, the mean, variance, skewness, and kurtosis of voxel subsets, to reflect regional texture variability related to amygdala cytoarchitecture. In contrast to qualitative approaches based on single features, such as investigations based on the detection of specific cell types (55), our pipeline captures several aspects of amygdala microstructure informing non-linear dimensionality reduction methods applied to a high-dimensional feature space. The resulting components U1 and U2 reflected complex combinations of central moments, with U1 being mainly scaled to mean intensities and U2 being associated to weighted combinations of the different moments. Crucially, we validated this coordinate space using openly available maps of amygdalar subdivisions from histological examinations performed by expert neuroanatomists (39). This approach complements previous work harnessing subnuclear parcellations of the amygdala derived from visual inspections of post mortem specimens (15) and deep-learning algorithms applied to *in vivo* MRI data (34,56,57) to contextualize variations in functional connectivity profiles within this region. Indeed, a significant advantage of our framework lies in the ease with which it may be applied to new datasets. For instance, our method overcomes the time-consuming nature and high level of expertise required for precise manual subnuclear segmentations. Furthermore, this approach circumvents the need for large datasets required to validate deep learning-based applications, a particular concern in the case of histological data which are often limited to few or even single specimens. However, it is important to note that both datasets analyzed in this work are limited by their small sample size (n=1 for BigBrain and n=10 for MICA-PNI). We speculate that the signal variations captured by U2 may be sensitive to artifacts or subject-specific sources of variance, potentially explaining why it was not consistent between subjects and modalities. Moving forward, this hypothesis could be assessed in future work via the analysis of larger histological and neuroimaging datasets to better track recurring features picked up by U2 or the association of these unique topographies with behavioral markers. Overall, the proposed framework lays the groundwork for future investigations of subcortical structure-function coupling by anchoring connectivity and task-related activations within measurements of regional microarchitecture.

The present works described a continuous coordinate space of amygdala subregional microstructure. However, this region has been previously described as consisting of three main subdivisions: LB, CM, and SF, each composed of smaller subnuclei with distinct connectivity patterns and functions (9,10,16,58). These subregions are largely conserved between humans and monkeys, reflecting their evolutionary relationship. However, there are still some considerable differences such as in the SF subregion, where its description in monkeys additionally contains the lateral olfactory tract (LOT) (59). Although qualitative histological accounts have indeed delineated multiple subunits within these general regions, the present work focuses on three subdivisions (16). To account for resolution disparities when translating our findings to in vivo MRI data. The LB subdivision includes the basomedial nucleus (Bm), basolateral nucleus (BL), lateral nucleus (LA) and paralaminar nucleus (PL). Moving medially, the CM subdivision includes the central (Ce) and medial nuclei (Me), while the SF subdivision includes the anterior amygdaloid area (AAA), amygdalohippocampal transition area (AHi), amygdalopiriform transition area (APir), and ventral cortical nucleus (VCo) (60). However, disagreement on the precise attribution of nuclei to broader subdivisions motivated our investigations of probabilistic subunits of the amygdala (15). The development of new tools to segment amygdala subnuclei *in vivo* opens opportunities for future work to further validate our framework at the precision of these nuclei within subjects (57). We selected UMAP for its potential to recover this nuclear architecture via the identification of discrete clusters of microstructural similarity within the amygdala. While these dimensions partially align with traditional concepts of arealization, they also provide a complementary, graded representation of amygdala microarchitecture. Although our framework leverages ultra-high resolution histological and myelin-sensitive MRI, the inherent spatial autocorrelation of feature intensities in these modalities may have emphasized the continuous signal variations we identify within the amygdala and hindered the discovery of discrete boundaries between known subdivisions. We address this limitation by benchmarking U1 and U2 distributions against validated probabilistic maps of major amygdala subdivisions (16,39), enabling us to recover the established biological validity of its nuclear organization. Furthermore, this approach allowed us to derive discrete clusters of maximal microstructural differentiation within the amygdala; these clusters served as seed regions for microstructurally-grounded and individualized investigations of the functional connectome embedding of amygdala subregions. Following an inferior-superior and medial-lateral axis of differentiation in both *post mortem* histology and myelin-sensitive *in vivo* MRI, this bipartite division is in line with previous work investigating the structural connectivity of the amygdala using diffusion-weighted imaging and probabilistic tractography (61) as well as functional connectivity from rsfMRI (62). Indeed, both modalities highlight the existence of two distinct clusters segregating amygdala connectivity to temporopolar and orbitofrontal cortices. These findings mirror macroscale associations seen in the neocortex between microstructure and connectivity, which emphasize close correspondence between the strength of interareal connectivity and microstructural similarity (63–65). In the case of the amygdala, our framework could thus recover these distinct anatomical pathways from a data-driven, texture-based analysis of microarchitectural information alone, supporting the potential of such contrasts to provide insights into the large-scale network embeddings of subcortical systems.

In line with this suggested association between amygdala microarchitecture and functional connectivity, we conclude our analyses by leveraging subject-specific representations of amygdala microstructure to map large-scale variations in its functional connectivity to the neocortex. We isolated and contrasted the highest and lowest 25% of U1 values for each participant to define an individualized bipartite parcellation of the amygdala. Qualitatively, we found that the subregion defined by the highest 25% of U1 values mainly overlapped with what is commonly defined as the superficial and centromedial subregions, whereas the lowest 25% U1 values subregion overlapped mostly with the laterobasal division. Interestingly, CM and SF characterized subregions showed significantly stronger functional connectivity to prefrontal structures. This finding aligns with previous work demonstrating unique affiliations between the CM subregion and anterior cingulate and frontal cortices (25,26), as well as between the SF subregion and the orbitofrontal cortex (17,24,30,66). Although these findings are promising, we also observe considerable overlap between functional connectivity networks of both our defined subregions. Indeed, the amygdala is a relatively small structure, leading to likely interconnectivity between its subregions and locally high signal autocorrelation. Functional connectivity and microstructure in the amygdala are certainly related, however previous work suggests they do not perfectly overlap (10). In addition, this region is affected by relatively low signal-to-noise ratio (SNR), as is observed in broader temporobasal and mesiotemporal territories. Decoding of subregional functional connectivity results indicated possible dissociations in cognitive (*e.g.,* memory) and affective *(e.g.,* emotional face processing) functions of the amygdala, echoing previous accounts of this region’s functional specialization and subregional segregation of associative processing of emotional stimuli. Notably, these findings link the functional connectivity profile of a subregion partially co-localizing with LB to emotional face processing. The LB subregion has been previously linked to associative processing related to the integration of sensory information (10,24,27–29), which is consistent with the association with visual emotional information processing identified in the present work. For the right amygdala, dissociation in functional connectivity patterns were more subtle, leading to overall similar functional decoding across the two clusters **(Figure S2)**. Overall, our findings suggest that this microstructurally-grounded delineation of U1 subregions could capture dissociations in their respective functional associations and potentially with fear-related processes. These results echo previous chemoarchitectural descriptions of the amygdala involving the 5-HT receptor, which has been closely associated with fear responses in mice and humans (31,67). Indeed, this receptor is expressed in lower densities in the CM region, overlapping with our U1 subregions that show lower connectivity to regions involved in fear-related responses. The present work thus offers an important step towards a more integrated account of the amygdala’s microstructural composition and functional organization.

By harnessing an openly available arsenal of tools and methods from histology, radiomics, and neuroinformatics, we define a comprehensive framework enhancing our understanding of individual differences in amygdala organization. Our findings contribute to a growing body of research emphasizing the importance of integrating structural and functional data to elucidate the complex roles of the amygdala in both health and disease. This multimodal, multiscale, and subject-specific approach not only advances our knowledge of the amygdala’s microstructural and functional intricacies but also offers a valuable resource for future studies exploring subcortical structures and their implications in various neurological and psychiatric conditions. This integrated perspective is essential for developing more precise and personalized interventions for disorders associated with amygdala dysfunction.

## METHODS

### Histological data acquisition and pre-processing

Cell-body-staining intensity of the amygdala was obtained from the BigBrain dataset (42). BigBrain is an ultra-high–resolution Merker-stained 3D volumetric histological reconstruction of a *post mortem* human brain from a 65-year-old male, made available on the open-access BigBrain repository (<u>bigbrain.loris.ca</u>). The *post mortem* brain was paraffin-embedded, coronally sliced into 7,400 20-μm sections, silver-stained for cell bodies (9), and digitized. As such, imagem sections, silver-stained for cell bodies (9), and digitized. As such, image intensity values in this dataset provide direct measurements of brain cytoarchitecture. Following manual inspection for artifacts, automatic repair procedures were applied, involving nonlinear alignment to a *post mortem* MRI, intensity normalization, and block averaging (68). 3D reconstruction was implemented with a successive coarse-to-fine hierarchical procedure. All main analyses were performed using the 100μm sections, silver-stained for cell bodies (9), and digitized. As such, imagem isovoxel resolution dataset.

### Amygdala segmentation and subdivision mapping

The left and right amygdalae were isolated from the BigBrain volume using an existing manual segmentation of left and right subcortical structures (45). This segmentation was warped to BigBrain histological space from a standard template space (ICBM2009b symmetric (69)) using openly available co-registration strategies aggregated in the BigBrainWarp toolbox (45,46). Notably, these approaches were optimized to improve the alignment of subcortical structures (45). After registering the subcortical atlas to histological space and resampling the segmentation to an isovoxel resolution of 100μm sections, silver-stained for cell bodies (9), and digitized. As such, imagem, we generated unique binary masks isolating the left and right amygdalae and performed manual corrections on each mask (*i.e.,* improving smoothness and continuity of the mask borders), and eroded the mask by five voxels to provide a conservative estimate of regional borders. All main analyses were performed on left hemisphere data only, while the right hemisphere served as a validation dataset (**Supplementary Figure S1**).

To contextualize our data-driven histological mapping of the amygdala (see below), we leveraged openly available probabilistic maps of amygdala subnuclei derived from visual inspections of *post mortem* tissue specimens performed by expert neuroanatomists (15,39). The borders of amygdala subdivisions were traced in 10 *post mortem* brains. Following 3D reconstruction and alignment of each *post mortem* brain to a common template space, voxel-wise probabilistic maps for each subregion were computed by quantifying the consistency of label assignments across the 10 donor brains. For the present work, all available probabilistic maps of the amygdala, including large subdivision groups encompassing multiple amygdala subnuclei (*i.e.,* CM, LB, and SF subdivisions) were accessed from the EBrains repository (v8.2) (39). Regional probabilistic maps were warped from ICBM2009c asymmetric space to the ICBM2009b symmetric template using the SyN algorithm implemented in the Advanced Normalization Tools software (ANTs) (70). Each subregional probabilistic map was subsequently warped to BigBrain histological space (45,46) and was resampled to an isovoxel resolution of 100μm sections, silver-stained for cell bodies (9), and digitized. As such, imagem. We then generated a maximum probability map to parcellate the amygdala into its subdivisions by retaining the voxels with the highest 5% probability values of belonging to each subdivision.

### Histological feature extraction

We built a histological feature bank of amygdala cell-body-staining using methods from the field of radiomics (71). Our approach for feature selection was also inspired by quantitative cytoarchitectural analyses developed in foundational neuroanatomical studies (43), involving the parameterization of intensity profiles with four central moments to characterize regional cytoarchitecture across the neocortex. In the present work, we computed these same four central moments (*i.e.,* mean, variance, skewness, and kurtosis) of cell body staining intensities in the amygdala to characterize intensity differences across this region. We used pyRadiomics v3.0.1 (47) to compute voxel-based maps for each of the selected first-order features, varying the size of 3D-feature extraction to a voxel neighbourhood of 500μm sections, silver-stained for cell bodies (9), and digitized. As such, imagem to 2100μm sections, silver-stained for cell bodies (9), and digitized. As such, imagem (in 400μm sections, silver-stained for cell bodies (9), and digitized. As such, imagem increments). Outlier values (> 1 standard deviation from the mean intensity value) were excluded from moment calculations at each kernel size. This resulted in 20 distinct feature maps, capturing variations in intensity distributions within the amygdala at finer and coarser scales. These feature maps were normalized by z-scoring each feature at each kernel size.

### Dimensionality reduction of histological features

To capture and visualize the underlying structure of amygdala cytoarchitecture, we applied UMAP to our normalized histological feature bank (48). This algorithm was selected over other compression approaches for its scalability in the analysis of large datasets, as well as its ability to preserve both local and global data structure (72). Two UMAP hyperparameters controlling the size of the local neighbourhood as well as the local density of data points were kept at their default settings (n_neighbours_=15, dist_min_=0.1). The resulting low-dimensional embedding of higher-order histological features was contextualized in relation to the amygdala’s *x*, *y*, and *z* voxel coordinate space, and validated against previously described maximum probability map of the amygdala.

### In vivo MRI data acquisition

After establishing this framework with *post mortem* histological data, we assessed its generalizability to *in vivo*, myelin-sensitive MRI contrasts. We capitalized on quantitative T1 (qT1) relaxometry data collected in 10 participants at a field strength of 7 Tesla. This sequence has been shown to be sensitive to cortical myeloarchitecture (73–75), and could thus offer complementary insights into the microstructural organization of the amygdala. Our cohort of 10 adult participants (5 men, mean ± SD age = 27.3 ± 5.71 years) were all healthy, with no history of neurological or psychiatric conditions (76).

Scans were acquired at the McConnell Brain Imaging Centre (BIC) of the Montreal Neurological Institute and Hospital on a 7T Terra Siemens Magnetom scanner equipped with a 32-receive and 8-transmit channel head coil. Synthetic T1-weighted (UNI) and quantitative T1 relaxometry (qT1) data were acquired using a 3D-MP2RAGE sequence (0.5 mm isovoxels, 320 sagittal slices, TR=5170 ms, TE=2.44 ms, TI1=1000 ms, TI2=3200 ms, flip angle_1_=4°, iPAT=3, partial Fourier=6/8 flip angle_2_=4°, FOV=260×260 mm^2^). We combined two inversion images to minimize sensitivity to B1 inhomogeneities and optimize reliability (77,78). rsfMRI scans were acquired using a multi-echo, 2D echo-planar imaging sequence (1.9mm isovoxels, 75 slices oriented to AT-PC-31 degrees, TR=1690 ms, TE1=10.8 ms, TE2=27.3 ms, TE3=43.8 ms, flip angle=67°, multiband factor=3). The rsfMRI scan lasted ∼6 minutes, and participants were instructed to fixate a cross displayed in the center of the screen, to clear their mind, and not fall asleep.

### Multimodal MRI processing and analysis

*a) Anatomical segmentation and co-registration*. Image processing leading to the extraction of cortical and subcortical features and their registration to surface templates was performed via *micapipe v0.2.3*, an open multimodal MRI processing and data fusion pipeline (<u>github.com/MICA-MNI/micapipe/</u>) (79). The amygdala was automatically segmented on the T1w images with volBrain v3, in every subject (51). Cortical surface models were generated from MP2RAGE-derived UNI images using FastSurfer 2.0.0 (80,81). Surface extractions were inspected and corrected for segmentation errors via placement of manual edits. Native-surface space cortical features were registered to the fs-LR template surface using workbench tools (82).

*b) qT1 image processing and analysis*. Automated, individual-specific segmentations of the amygdala were obtained with VolBrain, and were manually inspected and corrected prior to image processing. In a similar fashion to the histological data, a feature bank was rendered from the same central moments, although kernel sizes varied from size 1-5, rather than 2-10 in order to take into account the difference in image resolution between the two datasets. Thus, voxel neighborhoods ranged from 1500μm sections, silver-stained for cell bodies (9), and digitized. As such, imagem to 5500μm sections, silver-stained for cell bodies (9), and digitized. As such, imagem (in 1000μm sections, silver-stained for cell bodies (9), and digitized. As such, imagem increments). This resulted in 20 distinct feature maps, capturing variations in intensity distributions within the amygdala at finer and coarser scales. The new feature bank was again submitted to UMAP for dimensionality reduction, independently for all 10 participants. This procedure generated a 2-dimensional feature space unique to each participant recapitulating the organization of the amygdala. We then plotted the values generated in our UMAP space onto their original coordinate points inside the amygdala in the original qT1 space of each subject for further visualization and analysis.

To finally compare subject-specific *in vivo* MRI findings to our previously defined histological components, we first co-registered each subject’s qT1 scan to the ICBM152 template and applied the resulting transform to each participant’s U1 map. We applied existing transformations to bring the histological-space U1 to the same template space (45,46). This allowed us to contrast the UMAP components of all 10 subjects and BigBrain, along *x, y,* and *z* coordinate axes.

*c) rsfMRI image processing and analysis.* Processing employed micapipe v.0.2.3 (79), which combines functions from AFNI (83), FastSurfer (80,81,84), workbench command (85), and FSL

(86). Images were reoriented, as well as motion and distortion corrected. Motion correction was performed by registering each timepoint volume to the mean volume across timepoints, while distortion correction utilized main phase and reverse phase encoded field maps. Leveraging our multi-echo acquisition protocol, nuisance variable signals were removed with tedana (87). Volumetric timeseries were averaged for registration to native FastSurfer space using boundary-based registration (88), and mapped to individual surfaces using trilinear interpolation. Cortical timeseries were mapped to the hemisphere-matched fs-LR template using workbench tools then spatially smoothed with a 10mm Gaussian kernel. Surface-and template-mapped cortical timeseries were corrected for motion spikes using linear regression of motion outliers provided by FSL. Functional and anatomical spaces were co-registered using label-based affine registration (89) and the SyN algorithm available in ANTs (90).

### Functional network mapping of amygdala microstructural subregions

We averaged the rsfMRI timeseries within amygdala subregions defined by the highest and lowest 25% of values in each participant’s own qT1-derived U1 component. Functional connectivity of each amygdala subregion to the rest of the cortex was determined using Pearson correlation between the timeseries of each amygdala subregion and each cortical vertex. Resulting correlation coefficients underwent Fisher R-to-Z transformation to increase the normality of the distribution of functional connectivity values.

A mixed effects model implemented with the BrainStat toolbox (91) assessed differences between the cortical connectivity profiles of each U1 subregion, while considering age and sex and fixed effects and subject identity as a random effect. We corrected findings for family-wise errors (FWE) using random field theory (p_FWE_ < 0.05; cluster-defining threshold (CDT) = 0.01). We further contextualized differences in each connectivity map using decoding functions from NeuroSynth (52) that are made available via BrainStat (91), by contrasting the overall connectivity profile of the amygdala to those of each of its microstructurally-defined subregions. First, we divided subregional amygdala-cortical connectivity profiles into seven established functional network communities (19) and contrasted their average connectivity strength across each network. Meta-analytic functional decoding of the connectivity patterns of both amygdala subregions and the whole amygdala also highlighted different cognitive affiliations of each seed. Spatial correlations between the amygdala’s cortical connectivity profile and spatial activation maps associated with each term allowed us to retain ten terms associated with different cognitive domains. We then compared the association between the activation patterns associated with each of the retained terms and the cortical connectivity profiles of each amygdala U1 subregion.

### Data and code availability

Analysis notebooks related to this project are available on GitHub (<u>github.com/MICA-</u> <u>MNI/micaopen/AmygdalaUMAP</u>). BigBrain data are available on (<u>osf.io/xkqb3/</u>), 7T data are available on the Open Science Framework <u>(osf.io/mhq3f/</u>).

## ACKNOWLEDGEMENTS

H.A. acknowledges funding from the Fonds de la Recherche du Québec – Nature et Technologie (FRQNT) Master’s Training Scholarship. D.G.C. acknowledges support from FRQ-Sante and Savoy Foundation. J.R. acknowledges support from CIHR. B.C.B. acknowledges support from NSERC, CIHR, SickKids Foundation, BrainCanada, Future Leaders Research Grant, Helmholtz International BigBrain Analytics and Learning Laboratory (HIBALL), Healthy Brains and Healthy Lives, FRQS, and the Canada Research Chairs program.

