## Supplementary Materials for "From histology to macroscale function in the human amygdala"

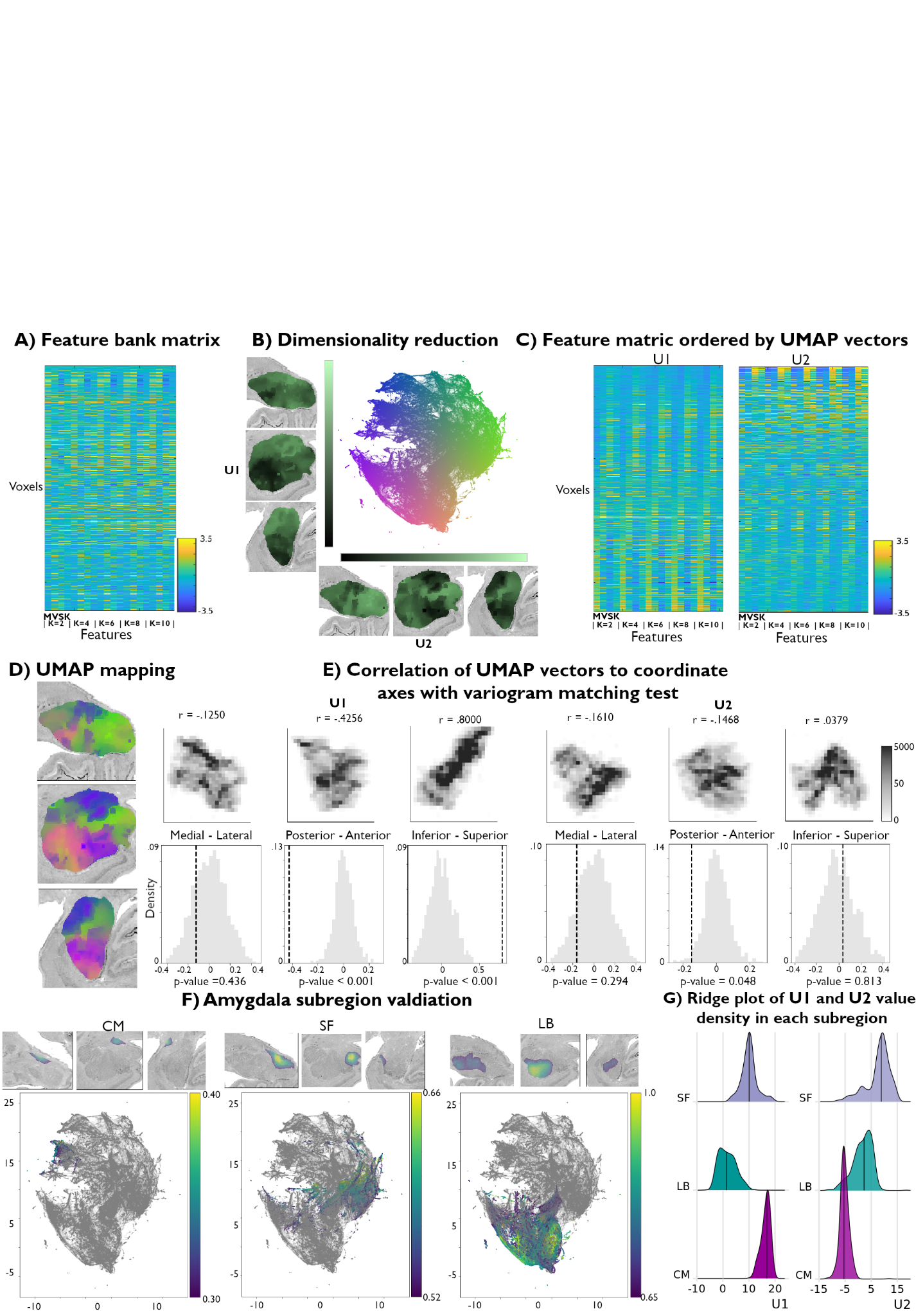

**Supplemental Figure S1. Replication of data-driven histological mapping for the right amygdala. (A)** Matrix representation of the normalized feature bank. **(B)** We applied UMAP to this feature bank to derive a low dimensional embedding of amygdala cytoarchitecture, defining a 2-dimensional coordinate space of amygdala cytoarchitecture (scatter plot, middle). Colors of the scatter plot represent proximity to axis limits. **(C)** Reordering the feature bank according to each eigenvector (U1 and U2) highlights the underlying variance in each feature captured by UMAP. **(D)** Coloring each amygdala voxel according to its corresponding location in the UMAP embedding space partially recovered its anatomical organization. **(E)** U1 and U2 were correlated to the three axes and a variogram matching test was done for each axis to assess the statistical significance of each correlation. **(F)** Coloring the embedding space with openly available probabilistic map labels of the three main amygdala subregions could recover the fully data-driven structure uncovered by UMAP. **(G)** Ridge plots of the probability values per subregions also illustrate a characterization of the subregions in U1.

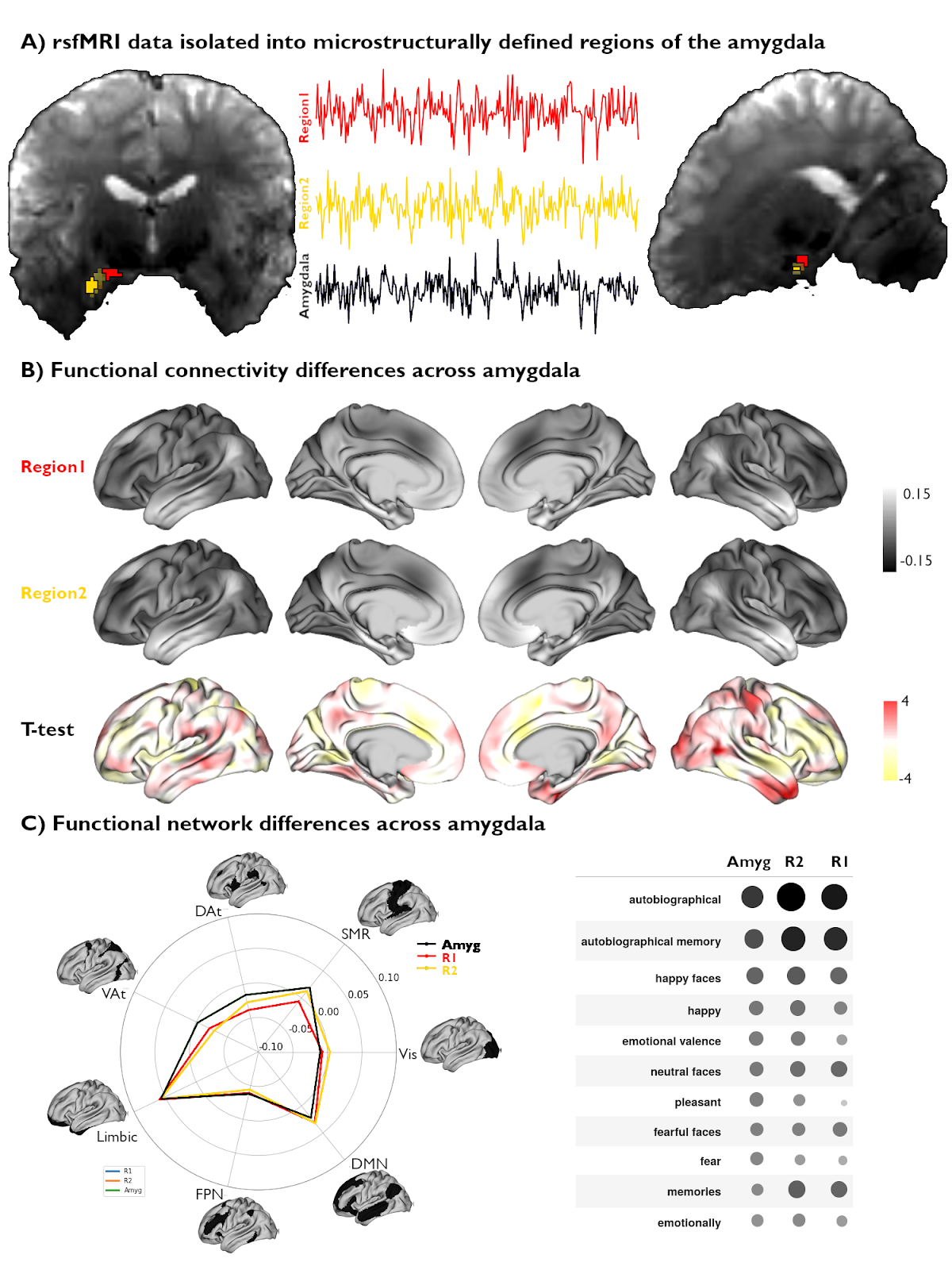

**Figure S2. Functional network mapping of amygdala microstructural subregions. (A)** We isolated the rsfMRI timeseries of two amygdala subregions (R1, R2), defined from subject-specific U1 topography, as well as the whole amygdala. (B) We computed the functional connectivity of both amygdala subregions, and projected resulting correlations to the cortex. Differences in connectivity patterns between subregions are shown as t-values across the cortex. (C) Left: Activation patterns illustrated in (B, top) were averaged within intrinsic functional communities defined by Yeo, Krienen, et al. (2011) to highlight differences in connectivity of each amygdala subregion in each network. Right: Meta-analytic decoding of functional connectivity patterns of both amygdala subregions and the whole amygdala showed similar functional affiliations of analyzed connectivity patterns.

**Supplementary table S1.1** Correlation values between UMAP parameter U1 and features.

| Kernel size: | 2 | 4 | 6 | 8 | 10 |
| --- | --- | --- | --- | --- | --- |
| Mean | 0.850 | 0.909 | 0.938 | 0.954 | 0.962 |
| Variance | -0.010 | 0.045 | 0.081 | 0.104 | 0.118 |
| Skewness | -0.238 | -0.223 | -0.205 | -0.195 | -0.196 |
| Kurtosis | 0.311 | 0.369 | 0.389 | 0.401 | 0.408 |

**Supplementary table S1.2** Correlation values between UMAP parameter U2 and features.

| Kernel size: | 2 | 4 | 6 | 8 | 10 |
| --- | --- | --- | --- | --- | --- |
| Mean | -0.011 | -0.021 | -0.029 | -0.035 | -0.038 |
| Variance | 0.399 | 0.493 | 0.537 | 0.555 | 0.555 |
| Skewness | 0.396 | 0.586 | 0.664 | 0.693 | 0.684 |
| Kurtosis | 0.260 | 0.441 | 0.503 | 0.509 | 0.481 |

**Supplementary table S2.1** U1 correlations (r) with all 3 axes across all subjects and BigBrain **(*p_null_* < 0.05).**

|  | S1 | S2 | S3 | S4 | S5 | S6 | S7 | S8 | S9 | S10 | BB |
| --- | --- | --- | --- | --- | --- | --- | --- | --- | --- | --- | --- |
| I-F | **0.71** | **0.77** | **0.65** | **0.54** | **0.69** | **0.71** | **0.70** | **0.57** | **0.86** | **0.68** | **0.82** |
| P-A | 0.11 | **0.31** | 0.01 | 0.06 | 0.02 | 0.02 | 0.18 | 0.15 | 0.10 | 0.10 | **0.51** |
| M-L | **0.77** | **0.47** | **0.59** | **0.66** | **0.61** | **0.80** | **0.77** | **0.67** | **0.68** | **0.85** | 0.15 |

**Supplementary table S2.2** U2 correlations (r) with all 3 axes across all subjects and BigBrain **(*p_null_* < 0.05).**

|  | S1 | S2 | S3 | S4 | S5 | S6 | S7 | S8 | S9 | S10 | BB |
| --- | --- | --- | --- | --- | --- | --- | --- | --- | --- | --- | --- |
| I-F | **0.56** | 0.11 | 0.29 | **0.52** | **0.35** | **0.58** | 0.15 | 0.13 | 0.13 | 0.14 | 0.02 |
| P-A | 0.14 | 0.10 | **0.34** | **0.26** | 0.16 | **0.47** | **0.50** | **0.43** | **0.29** | **0.20** | **0.25** |
| M-L | 0.24 | 0.04 | 0.08 | 0.21 | **0.27** | 0.25 | **0.58** | **0.55** | 0.19 | **0.70** | 0.22 |
